# Massively parallel characterization of insulator activity across the genome

**DOI:** 10.1101/2022.11.29.518444

**Authors:** Clarice KY Hong, Alyssa A Erickson, Jie Li, Arnold J Federico, Barak A Cohen

## Abstract

Insulators are *cis*-regulatory sequences (CRSs) that can block enhancers from activating target promoters or act as barriers to block the spread of heterochromatin. Their name derives from their ability to ‘insulate’ transgenes from genomic position effects, an important function in gene therapy and biotechnology applications that require high levels of sustained transgene expression. In theory, flanking transgenes with insulators protects them from position effects, but in practice, efforts to insulate transgenes meet with mixed success because the contextual requirements for insulator function in the genome are not well understood. A key question is whether insulators are modular elements that can function anywhere in the genome or whether they are adapted to function only in certain genomic locations. To distinguish between these two possibilities we developed MPIRE (<u>M</u>assively <u>P</u>arallel Integrated <u>R</u>egulatory <u>E</u>lements) and used it to measure the effects of three insulators (A2, cHS4, ALOXE3) and their mutants at thousands of locations across the genome. Our results show that each insulator functions in only a small number of genomic locations, and that insulator function depends on the sequence motifs that comprise each insulator. All three insulators can block enhancers in the genome, but specificity arises because each insulator blocks enhancers that are bound by different sets of transcription factors. All three insulators can block enhancers in the genome, but only ALOXE3 can act as a heterochromatin barrier. We conclude that insulator function is highly context dependent and that MPIRE is a robust and systematic method for revealing the context dependencies of insulators and other *cis*-regulatory elements across the genome.

## Introduction

A consistent model of insulator function has yet to emerge from one-at-a-time perturbations of specific insulator elements. The effects of perturbations to putative insulators on genome structure and gene expression appear to depend on the specific locus and regulatory sequences involved^1^. A proposed model of insulator activity states that context-specific effects arises from how insulators organize genome structure to facilitate or restrict access to *cis*-regulatory sequences^2^. Yet, genome-wide depletion of known insulator binding proteins CCCTC-binding factor (CTCF) and Cohesin leads to widespread changes in genome structure but have little impact on gene expression^3,4^, suggesting that insulators may not act solely through genome organization. It is therefore important to directly assess the impact of insulators on gene expression across diverse genomic contexts.

Two classes of insulator activities have been described. Barrier insulators halt the spread of flanking heterochromatin, while enhancer-blocking insulators prevent enhancers from activating their target promoters when placed between the enhancer and promoter^2^. Enhancer-blocking assays are typically performed with one enhancer-promoter pair on plasmids, but some insulators only block specific enhancer-promoter pairs^5–7^. In the genome, some insulators have been shown to protect transgenes from trapped enhancers at ectopic locations and transgene silencing^8–10^. However, not all enhancer-blocking insulators protect against position effects^11^, suggesting that enhancer-blocking activity on plasmids does not necessarily correlate with insulator function in the genome. Insulator activity is also highly dependent on genomic context^12–14^. These results suggest that the impact of insulators on gene expression will be specific for certain locations in the genome, but it remains unclear how widespread insulator activity is and what the context dependencies of different insulators are.

The sequence features that contribute to insulator activity are also not well understood. In mammalian cells, the most well-characterized protein with insulation activity is CTCF. Yet, in the extensively characterized cHS4 insulator element, CTCF is only required for enhancer-blocking activity on plasmids, but not for protection against position effects in the genome^15–17^ and CTCF is neither necessary nor sufficient to block the spread of repressive H3K27me3 heterochromatin^3,12,18,19^. RNA pol III transcribed elements, including short interspersed nuclear elements (SINEs) and tDNA sequences, can possess both enhancer-blocking and barrier activity^20– 22^, but the sequence requirements for RNA pol III insulator activity is unclear. Given our current understanding, we cannot reliably predict the effects of genetic variants in insulator sequences on gene expression. Systematic studies of insulator elements and their genetic variants will be required to define how the critical sequences that comprise insulator function across the genome.

A major challenge to studying insulators is that their functions are intimately tied to their locations in the genome. Methods to measure the activity of a reporter gene integrated at a large number of genomic locations currently rely on random integration^23–25^, so different insulators cannot be compared to each other at the same locations^14^. Direct comparisons at the same genomic location can be achieved with recombinase mediated cassette exchange (RCME), where researchers can integrate libraries of reporter genes into a small number of defined genomic locations^26,27^. Walters *et al*. used a similar approach to assay the ability of the cHS4 insulator to block HS2 enhancer mediated activation of the γ-globin promoter at three genomic locations in K562 cells and found that cHS4 activity was exquisitely context-specific. However, gaining the power to unravel the features of genomic environments that determine insulator function will require a technology that can assay different insulator variants in a large number of genomic locations. Moreover, to reliably compare different insulators, each insulator sequence must be assayed at the same set of locations in the genome.

To address this problem we developed <u>M</u>assively <u>P</u>arallel Integrated <u>R</u>egulatory <u>E</u>lements (MPIRE), a method that measures the effects of different *cis*-regulatory sequences at thousands of defined genomic locations simultaneously. Using MPIRE, we present the largest characterization of three insulator sequences (cHS4, A2 and ALOXE3) at more than 10,000 locations across the genome. Since insulator activity is thought to depend on either CTCF binding sites (cHS4 and A2) or B box motifs (ALOXE3), we also measured the effects of mutations in the respective motifs at the same genomic locations. We show that each insulator functions at specific and distinct locations in the genome, with each insulator blocking enhancers bound by unique sets of transcription factors. This insulator activity is largely dependent on CTCF or the B box motif respectively. While all insulators can block enhancers in the genome, only ALOXE3 can act as a heterochromatin barrier. Our results suggest that protection against position effects results from diverse, context-dependent functions of insulators.

## Results

### MPIRE measures CRS activity at defined locations genome-wide

The goal of MPIRE is to integrate reporter genes carrying different *cis*-regulatory sequences into large numbers of locations in parallel. The key reagents in MPIRE are pools of cells that carry large numbers of genome-integrated landing pads^26,28^. The landing pads are integrated randomly into K562 cells where they can serve as sites for recombination-mediated cassette exchange (Fig 1A). We use the Sleeping Beauty transposon system to deposit multiple landing pads per cell in an unbiased manner throughout the genome^29^. We selected φC31 and BxbI attP sites for recombination because they were previously shown to be highly efficient^30^, and each individual landing pad integration carries a unique genomic barcode (gBC). In total we generated 6 cell pools (Pools 1-6), with each pool containing between 2000-6000 uniquely mapped gBCs (Supplementary Fig 1A, Supplementary Table 1).

**Fig 1:**
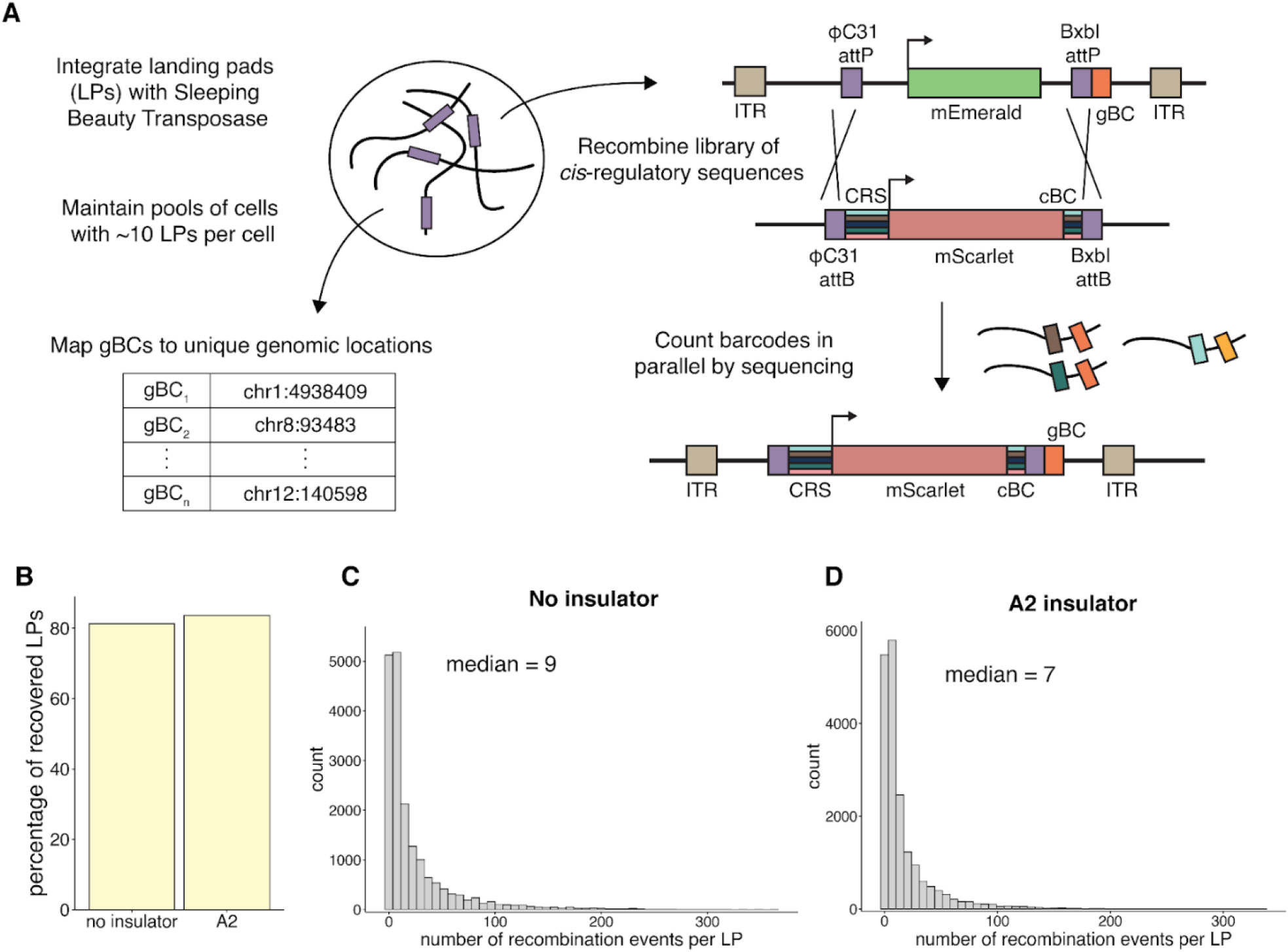
MPIRE is an efficient method for measuring CRS activity genome-wide. **(A)** Overview of MPIRE method. Pools of cells are generated by integrating landing pads driven by the Cytomegalovirus (CMV) promoter into the genome using SB transposase. The gBCs are mapped to a unique location in the genome and can be used to recombine CRSs for parallel measurements of *cis*-regulatory activity across the genome. gBC: genomic barcode. ITR: inverted terminal repeats. CRS: *cis*-regulatory sequence. cBC: CRS barcode. **(B)** Proportion of landing pads mapped to a unique location that were recovered after recombination with different CRS constructs. **(C, D)** Distribution of the number of recombination events per location for an uninsulated construct **(C)** and construct with the A2 insulator **(D)**.

Once the pools are generated, reporter genes carrying different insulators can be integrated into the landing pad pools simultaneously. Reporter genes driven by the hsp68 promoter with different insulator sequences are cloned between recombination sites for integration into landing pad pools (Fig 1A) and marked with different *cis*-regulatory library barcodes (cBCs) so that the assay can be multiplexed. The reporters are pooled and recombined into each landing pad pool. After recombination, the expressed mRNAs from each landing pad will contain two barcodes: a cBC indicating the identity of the insulator sequence and a gBC indicating the genomic location of the landing pad. We then sequence and count the number of DNA and RNA BC-pairs to calculate expression of each reporter at each location in the genome.

We first verified the utility of landing pad pools for MPIRE. For these analyses, we focused on Pool 1. Using qPCR, we estimated that each cell carries approximately 11 independent landing pads. The landing pads are well-distributed across the genome (Supplementary Fig 1B), consistent with previous observations that Sleeping Beauty integrates in an unbiased manner^29^. Expression from these landing pads span a large range of expression (>10^4^-fold) indicating that the integrations are sampling diverse genomic environments (Supplementary Fig 1C, Supplementary Table 2). To test whether different landing pads in the same cells might recombine with each other, we transfected cells with integrase only (no CRS library) and found that the landing pads have similar expression levels and mostly map back to the same locations (Supplementary Figs 1D and 1E). Thus, the landing pads in our pools represent diverse genomic environments and landing pads in the same cell do not interfere with each other.

We next evaluated the efficacy of large-scale recombination across locations in Pools 1 and 2. We measured the integration efficiency of two reporter genes, one containing only the minimal hsp68 promoter and one that also contains the A2 insulator^31^ to test whether the insulator sequence might impact recombination efficiencies (Fig 2A). We barcoded each reporter gene with a highly diverse library of random barcodes (rBCs) and transfected them independently into both pools of cells. Since the library contains ∼500,000 rBCs, each integration is likely associated with a unique rBC such that the number of rBCs per gBC represents the number of independent integrations at each location. We obtained barcodes from ∼80% of all mapped landing pads in pool 1 for both the no insulator and A2 constructs (Fig 1B). For each landing pad, we obtained a median of 9 independent integrations for the no insulator reporter and 7 independent integrations for the A2 reporter (Fig 1C & 1D) for a total of 18,492 and 19,031 integrations, respectively. These results demonstrate that MPIRE is a robust method for measuring *cis*-regulatory activity at a large number of defined locations across the genome.

**Fig 2:**
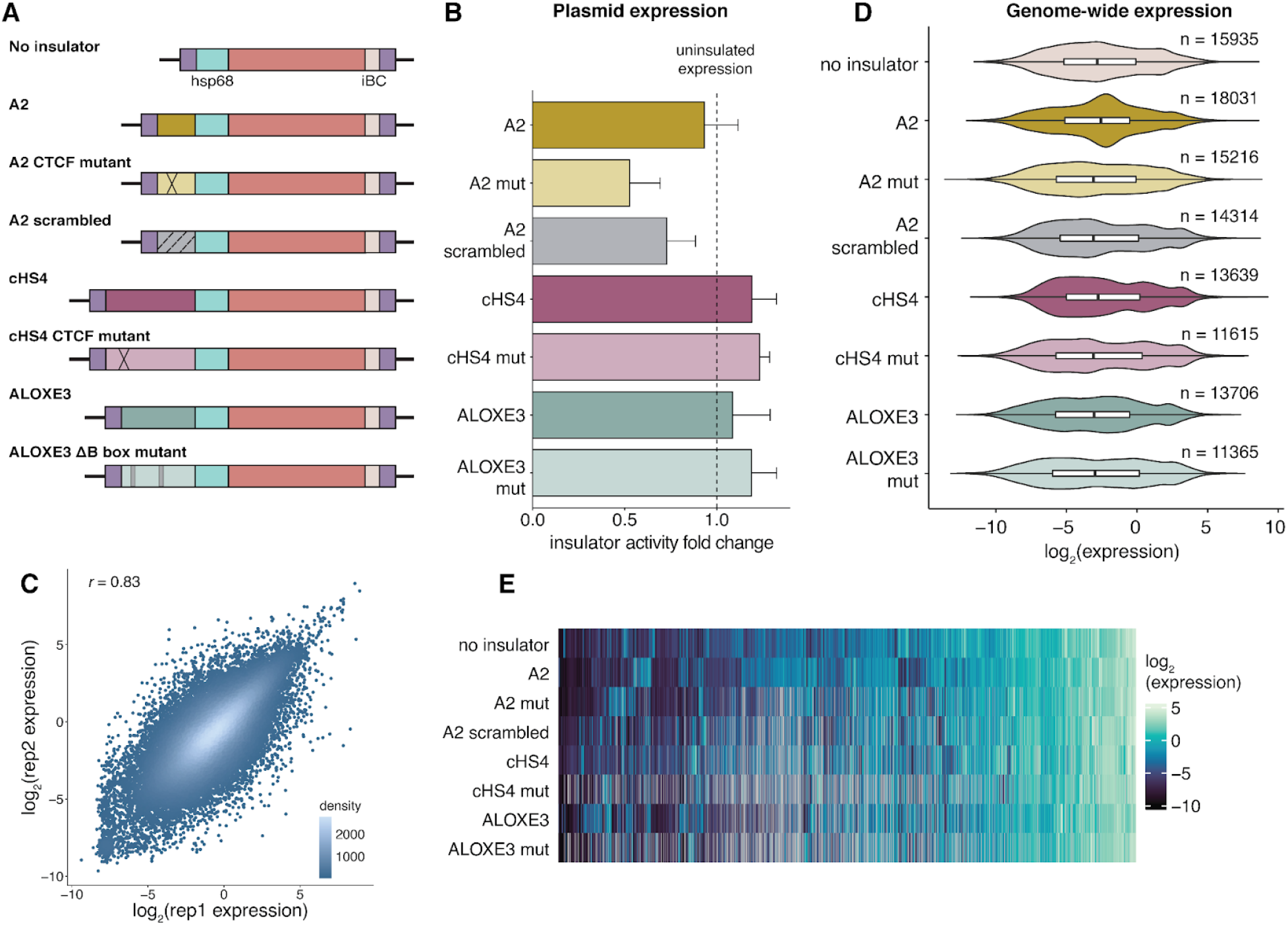
Insulators do not have global effects on gene expression. **(A)** Insulator constructs and mutants used in this experiment. **(B)** Insulator activity on plasmids measured by qPCR calculated as fold-change over the uninsulated construct (dotted line denotes fold-change = 1). Error bars represent the SEM from three biological replicates. **(C)** Reproducibility of measurements from two biological replicates. Correlation indicated is Pearson’s *r*. Color represents the density of neighboring points. **(D)** Expression distribution of each insulator across all locations. n indicates the number of locations measured for each construct. **(E)** Heatmap of expression across the genome, where each column represents a different location. Only locations where the ‘no insulator’ construct was measured is plotted to allow for comparisons against the uninsulated reporter.

### Measurement of insulator activity across the genome

We selected three previously characterized insulators to test by MPIRE (Fig 2A). cHS4 is the most well-characterized insulator and is derived from 5′ end of the chicken β-globin locus^9^. A2 was identified in a screen of endogenous CTCF-containing sequences for enhancer-blocking activity^31^, and ALOXE3 tDNA was discovered in a screen for tDNA sequences with insulator activity^21^. These studies showed that all three insulators function in canonical enhancer-blocking assays in K562 cells. The enhancer-blocking activity of cHS4 depends on its CTCF binding site^15,32^, while the enhancer-blocking activity of ALOXE3 depends on its two B box motifs, which recruit RNA polymerase III^33^. CTCF binds to A2 at 100% occupancy in the K562 genome, suggesting that CTCF is likely to be important for A2 activity^31^. To test the importance of binding motifs we mutated the CTCF binding sites in cHS4 and A2 (cHS4-mut and A2-mut, respectively) and deleted both instances of the B box motif in ALOXE3 (ALOXE3-mut) (Fig 2A). The cHS4-mut sequence was previously shown to abolish CTCF binding^32^, while the A2-mut sequence was designed to mutate the most nucleotides in the CTCF position-weight matrix. In addition, we scrambled the A2 insulator (A2-scrambled) to test the effects of a random DNA sequence that lacks identifiable *cis*-regulatory elements.

Before performing MPIRE with these insulators, we confirmed their enhancer-blocking activity in a plasmid-based assay (Supplementary Fig 2A). The A2-mut and ALOXE3-mut constructs had less enhancer-blocking activity than their wild-type counterparts in the plasmid assay, but not the cHS4-mut construct. We also measured whether any of the insulator sequences had *cis*-regulatory activity that was independent of their enhancer-blocking activity. In the absence of an enhancer, the insulators had minimal effects on reporter gene expression (Fig 2B), with the exception of the A2-mut, which had some independent repressive activity. However, this repressive activity was not recapitulated globally in the genome (see Fig 2D below). Thus, the insulators we tested do not directly influence promoter activity, but do have enhancer-blocking activity.

We performed MPIRE by pooling the insulator constructs and transfecting them into six pools of landing pad cells. In total, we measured expression from each insulator construct in 11,365-18,031 landing pad locations (Supplementary Table 3). The measurements are reproducible across biological replicates consisting of independent transfections (Fig 2C). The expression of the non-insulated construct was well correlated with the expression of the landing pads before recombination (Supplementary Fig 2B). These results suggest we were able to obtain accurate and reproducible measurements of multiple *cis*-regulatory sequences across thousands of genomic locations in parallel.

We first asked whether any of the insulators had widespread effects across the genome. In general, insulators did not have a large impact on the overall distribution of expression from locations across the genome (Fig 2D). Formally, this result could arise if insulators increased and decreased expression at different locations in a way that did not alter the overall distribution of expression. However, we find that expression largely depends on genomic location regardless of the identity of the insulator (Fig 2E), and that the expression of landing pads with and without insulators are well-correlated, with mutants slightly more correlated overall (Supplementary Figs 2C & 2D). These results are inconsistent with a model of insulators as modular elements that can function in a wide variety of genomic locations.

### Insulators function in a small subset of locations that are distinct from each other

While insulators do not appear to have global effects on expression across the genome, there are some locations where expression of the reporter gene with insulators is higher or lower than expected given its location in the genome (Fig 2E). We sought to identify the specific locations where insulators might be influencing expression. We first calculated fold-changes in expression for each insulated and non-insulated construct at each landing pad location in the genome. The number of landing pads with matched insulated and non-insulated measurements ranged from 10,944-14,390 for each insulator. A positive fold-change indicates a location where the insulator upregulates expression relative to the non-insulated construct, while a negative fold-change indicates that the insulator downregulates expression. We defined locations as ‘insulated’ if the insulator caused a greater than 2-fold change in expression (in either direction) with false discovery rate < 0.05 (Figs 3A-3C, Supplementary Figs 3A-3D, Supplementary Table 4). As expected from the lack of global insulator activity, we identified only a small number of genomic locations insulated by each insulator (ranging from 10-15%), with even fewer insulated by the mutant insulators (Fig 3D). All insulator and mutant constructs can lead to both upregulation and downregulation of expression, but the proportion of locations that are upregulated or downregulated varied by insulator (Fig 3E). This suggests that insulators may perform more than one function in a context- and insulator sequence-dependent manner. For the rest of this manuscript we refer to upregulated and downregulated locations as insulator-up (eg. A2-up) or insulator-down (eg. A2-down) respectively.

**Fig 3:**
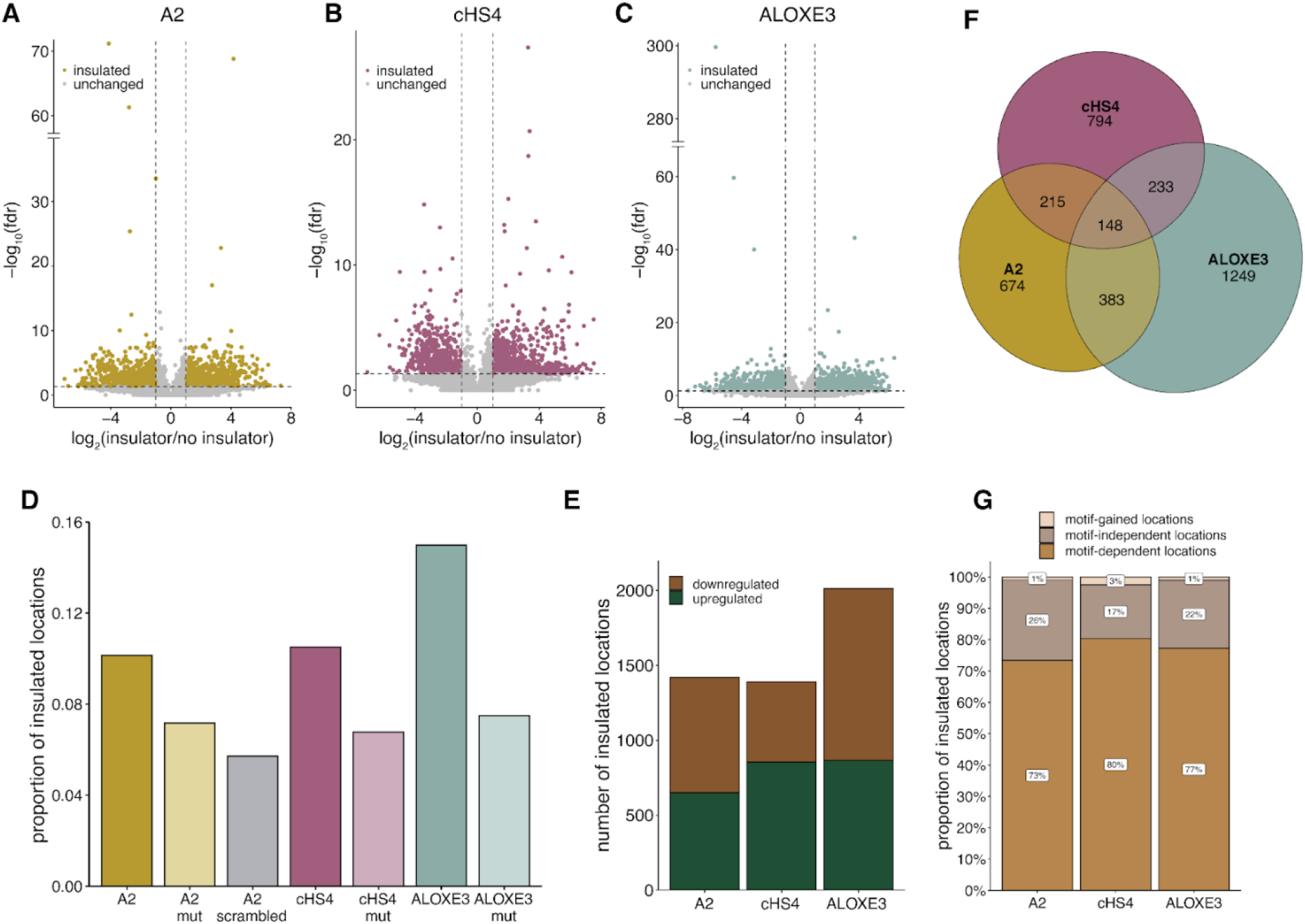
Insulators function in a small number of distinct locations in the genome. **(A, B, C)** Volcano plot of insulator effect across locations for A2 (n=14003) **(A)**, cHS4 (n=13223) **(B)** and ALOXE3 (n=13429) **(C)**. Colored points (insulated locations) represent locations where log_2_(insulator/no insulator) > 2 and fdr < 0.05. **(D)** Proportion of locations defined as ‘insulated’ for each construct. **(E)** Number of locations that are upregulated (log_2_(insulator/no insulator) > 0) or downregulated (log_2_(insulator/no insulator) < 0) by the insulator. **(F)** Venn diagram of ‘insulated’ locations shared between different insulator constructs. **(G)** Percentage of insulated locations that are motif-dependent, motif-independent, or motif-gained (defined in text) for each insulator.

The insulators could be insulating against common features in the genome, or each insulator may act at distinct locations. The overlap between insulated locations for the three insulators is small, with only 148 insulated locations shared between insulators (Fig 3F), suggesting that each insulator mostly functions at distinct genomic locations. However, this overlap is still greater than expected by chance (bootstrap *p*-value = 1×10^−4^), indicating that there might be some shared characteristics between insulated locations. The overlap is similar for both up- and downregulated locations (Supplementary Figs 3E, 3F), and most of the shared locations (92%) are insulated in the same direction by all three insulators (Supplementary Fig 3G). Because A2 and cHS4 both contain CTCF binding sites, we asked if there was more overlap between cHS4/A2 shared regions compared to ALOXE3, but the overlap between any two insulators is similar (Supplementary Fig 3H). These results suggest that each insulator requires a unique type of genomic environment to function and that CTCF is not sufficient to explain insulator activity.

We next asked if insulator activities depend on their respective CTCF or B-box motifs. The distribution of fold-changes of mutant versus wild-type insulators was very similar to the fold-changes of insulated versus uninsulated constructs, indicating that even a few nucleotide changes in the insulator sequence is sufficient to generate large changes in gene expression at the same location (Supplementary Figs 4A-4C). Using our definition of ‘insulated’ locations above, mutating an insulator can lead to three possible outcomes - the location could still be insulated in the same direction (motif-independent), insulated but in the opposite direction (motif-gained), or could no longer be insulated (motif-dependent). We find that 70-80% of the insulated locations are motif-dependent (Fig 3G), with the other locations being mostly motif-independent. The proportion of motif-dependent locations is also similar for both up- and downregulated locations (Supplementary Figs 4D & 4E), suggesting that the motifs are necessary for both classes of insulator activity. The A2-scrambled control also has higher numbers of motif-dependent locations compared to the A2-mut, suggesting that there might be other sequences in A2 that contribute to insulator activity in addition to the CTCF sites (Supplementary Fig 4F). Our results suggest that insulators only function in small numbers of locations in the genome and that their CTCF or B-box motifs are necessary for insulator activity.

### Insulators block specific enhancers to downregulate expression

We next focused on locations where the addition of an insulator leads to a decrease in gene expression. At these downregulated locations, insulators may be blocking the effects of nearby enhancers. Thus, we predicted that down-regulated locations are normally highly active genomic environments and should contain higher levels of active chromatin modifications. Consistent with this prediction, insulator-down locations are indeed highly enriched in active chromatin modifications, and depleted for H3K27me3, a marker of Polycomb-repressed heterochromatin^34^ (Fig 4A). Since the insulator is only present on the 5’ end of the reporter, we asked if there was more enrichment upstream compared to downstream of insulator-down locations, but found similar enrichment of signals both upstream and downstream (Supplementary Fig 5A-5B). We did not observe the same enrichment at insulator-down locations by the mutant constructs (Fig 4B), consistent with the idea that the mutations abrogate insulator activity. Thus, insulator-down locations are enriched for active histone modifications, supporting our model that insulators block enhancers in a motif-dependent fashion at these locations.

**Fig 4:**
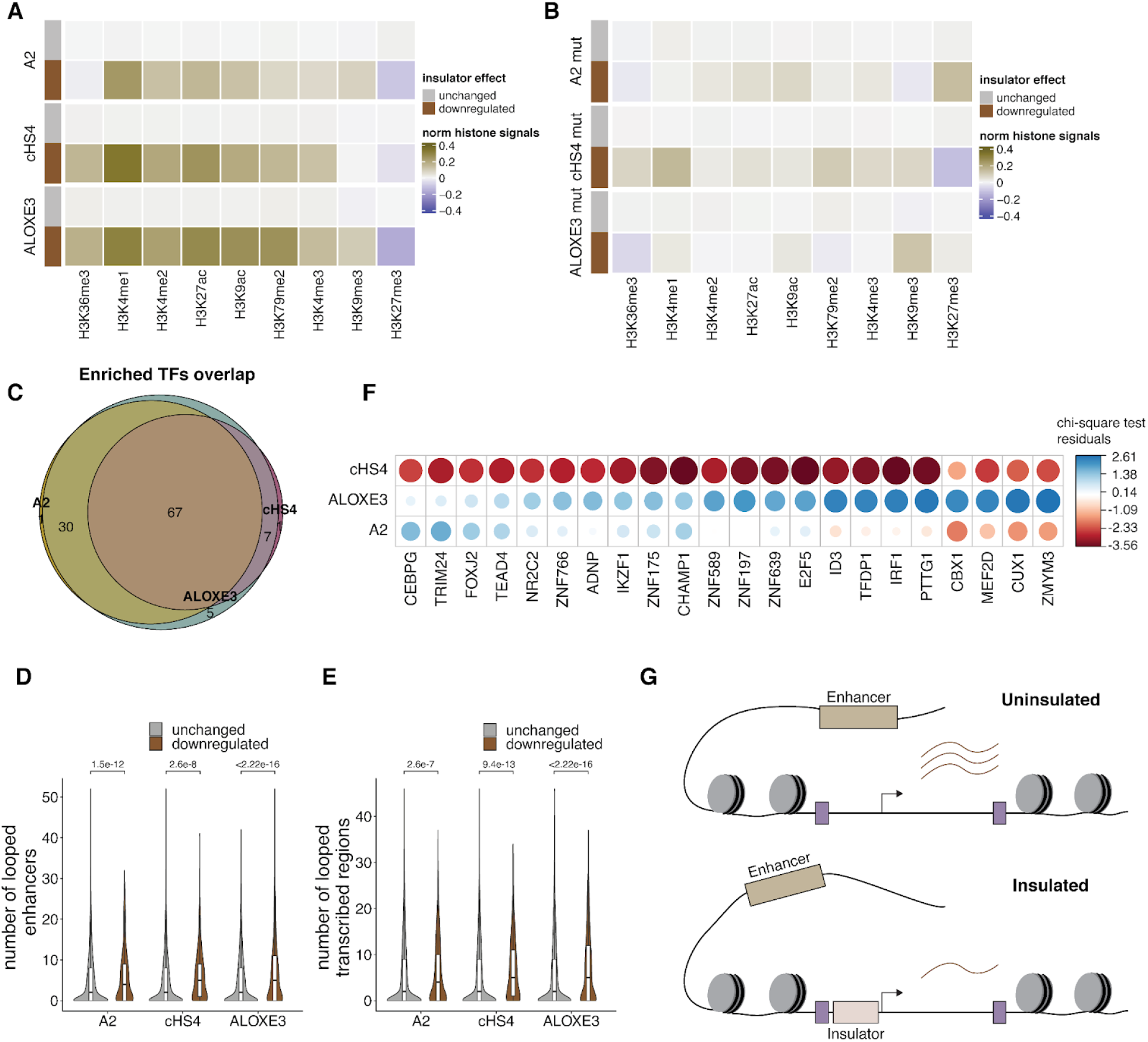
Insulators block specific enhancers in the genome. **(A)** Mean normalized histone modifications for insulator-unchanged and insulator-down locations. Histone signals are calculated as the mean of 10kb surrounding each location and standardized across locations for each modification. Insulator-down locations are enriched for ‘active’ histone modifications. **(B)** Mean normalized histone modifications for mutant-unchanged and mutant-down locations. Mutant-down locations are generally not enriched for any histone modifications. **(C)** Venn diagram of overlap in enriched TFs between insulator-down and insulator-unchanged locations. The number of TF binding peaks in the 10kb surrounding each location was used to calculate enrichment by chi-squared tests. Only TFs that are significantly differentially bound (*p* < 0.05 afterBenjamini & Hochberg (BH) correction) are included in this plot. **(D)** Violin plot of number of looped enhancers per location for insulator-unchanged vs insulator-down locations. **(E)** Violin plot of number of looped transcribed regions per location for insulator-unchanged vs insulator-down locations. *p*-values (two-sided Wilcoxon test) are shown above each plot. **(F)** For all insulators, enhancers that are looped that the insulator-down locations were compared for differences in TF binding measured by ChIP (chi-squared test). The residuals of TFs that are significantly different (*p* < 0.05) after BH correction was plotted. **(G)** Model for how insulators downregulate expression at specific locations. All insulators are located in open chromatin regions as depicted by the nucleosomes. The insulator blocks enhancers interacting with the location from activating expression, leading to downregulation of expression.

Although the same histone modifications were enriched at the insulator-down locations for all three insulators, the different insulators still functioned at distinct genomic locations (Supplementary Fig 3A). This observation suggests that the insulators are not simply protecting the reporter gene against active chromatin modifications. Instead, each insulator may block distinct transcription factors (TFs) that are present in different genomic environments. In this model, insulators may block TFs that directly occupy the locations surrounding the landing pad, or the insulators may block TFs that bind distal enhancers which loop to these locations.

We first addressed whether TFs that directly occupy insulator-down locations mediate the specificity of different insulators. This model predicts that different TFs will be bound at locations that are downregulated by the different insulators. In contrast to this prediction, when comparing TF binding in the 10kb surrounding insulator-unchanged and insulator-down locations, the TFs that are enriched in insulator-down locations are similar for each insulator (Fig 4C). This suggests that the TFs that occupy insulator-down locations do not mediate the specificity of different insulators.

Instead, we hypothesized specificity may arise because each insulator blocks different TFs that bind the distal enhancers that loop to insulator-down locations. To address this hypothesis we identified putative enhancers for each insulator-down location from Hi-C data and overlapped these predictions with genomic annotations from chromHMM^35^. Insulator-down locations tend to interact with more enhancers and transcribed regions than insulator-unchanged regions (Fig 4D & 4E), consistent with the idea that insulator-down locations are in more active genomic regions. This difference was reduced in cHS4-mut-down locations and not present in A2-mut-down and ALOXE3-mut-down locations (Supplementary Figs 5C & 5D). Similarly, there were more ATAC-seq peaks in the 10kb surrounding insulator-down locations compared to insulator-unchanged locations (Supplementary Fig 5E). We then compared TF binding between enhancers looped to each of the insulator-down locations and identified TFs and motifs that are differentially enriched in each group of enhancers (Fig 4F, Supplementary Fig 5F). These results support the hypothesis that each insulator blocks a different set of enhancer-bound TFs. The general depletion of TF motifs in cHS4-down enhancers could be due to the fact that cHS4 is a longer sequence that contains more TF binding sites than the other two insulators and may therefore interact with a more diverse set of TFs (Supplementary Fig 5G). The lack of overlap in enriched TFs between A2 and cHS4, both of which bind to CTCF, also suggests that the specificity is not purely driven by CTCF. Taken together, insulators appear to block endogenous enhancers from influencing gene expression, and the specificity of insulator activity seems to be at least partially explained by the suite of TFs that are present in the enhancers interacting with each location (Fig 4G).

### ALOXE3, but not cHS4 or A2, acts as a heterochromatin barrier

Insulator-up locations are upregulated by insulators and are candidates for locations where insulators block the spread of repressive chromatin. This model predicts that insulator-up locations will be enriched for markers of heterochromatin. In contrast to the insulator-down locations, the chromatin landscape of insulator-up regions is not the same for the three different insulators (Fig 5A). ALOXE3-up locations are depleted of active histone modifications and enriched for H3K27me3, suggesting that ALOXE3 can block repressive histone modifications as expected for a heterochromatin barrier. Only ALOXE3-up locations are close to heterochromatin (Supplementary Fig 6A) and enriched for looping with repressed (Polycomb) regions (Supplementary Fig 6B). Similarly, ALOXE3 constructs are repressed at the least number of locations across the genome, suggesting that ALOXE3 can efficiently block heterochromatin silencing at many locations across the genome (Supplementary Fig 6C). Notably, only H3K27me3, an indicator of facultative heterochromatin, is enriched at ALOXE3-up locations. H3K9me3, which is associated with constitutive heterochromatin, is not enriched in ALOXE3-up locations, consistent with the idea that only facultative heterochromatin remains accessible to TF binding, allowing for upregulation once ALOXE3 is present to block H3K27me3 spreading^36^. This result is also consistent with the fact that ALOXE3 was derived from the boundary of a H3K27me3 domain in K562 cells^21^. In contrast to ALOXE3-up locations, cHS4-up locations are not enriched for any histone modifications, and A2-up locations are enriched for similar chromatin marks as A2-down locations, with a stronger enrichment for H3K36me3 (Fig 5A). From these results we propose that only ALOXE3 acts as a barrier against H3K27me3 spreading, while cHS4 and A2 upregulate expression by other mechanisms.

**Fig. 5:**
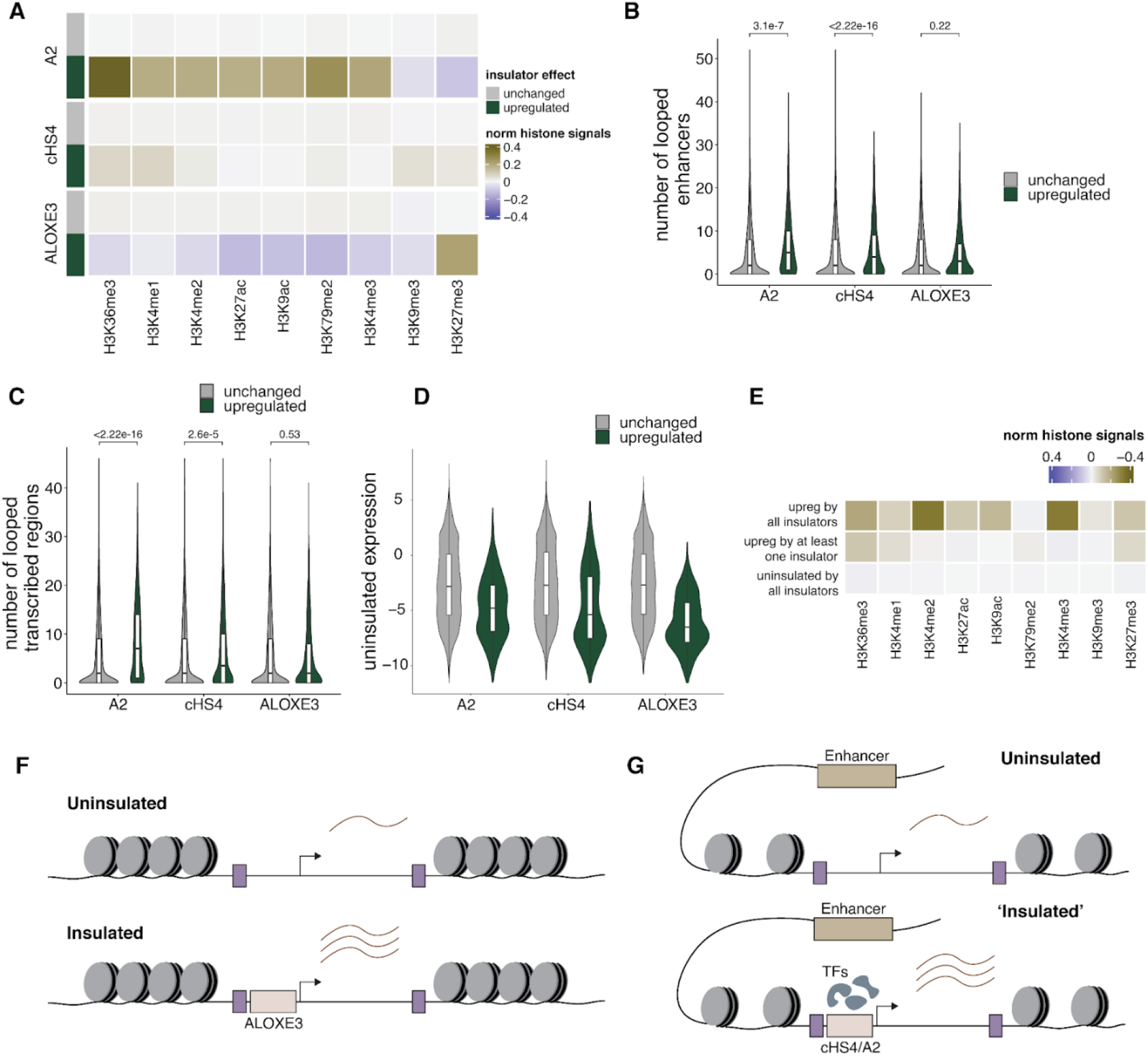
ALOXE3 acts as a heterochromatin barrier, while A2 and cHS4 act as enhancers at ‘primed’ locations. **(A)** Mean normalized histone modifications for insulator-unchanged and insulator-up locations. Histone signals are calculated as the mean of 10kb surrounding each location and standardized across locations for each modification. **(B)** Violin plot of number of looped enhancers per location for insulator-unchanged vs insulator-up locations. **(C)** Violin plot of number of looped transcribed regions per location for insulator-unchanged vs insulator-up locations. *p*-values (two-sided Wilcoxon test) are shown above each plot. **(D)** Expression distribution of the uninsulated reporter at insulator-unchanged vs insulator-up locations. **(E)** Mean normalized histone modifications for locations upregulated by all insulators, locations upregulated by only one or two locations and uninsulated locations. **(F)** Model for barrier activity at ALOXE3-up locations. ALOXE3 blocks heterochromatin from silencing the gene. **(G)** Model for cHS4/A2-up locations. Instead of acting like an insulator, we propose that cHS4 and A2 act like enhancers in ‘primed’ genomic environments to increase expression.

Our working model predicts that repressive H3K27me3 marks should decrease at insulated locations when ALOXE3 is present. We attempted to test this prediction computationally using Enformer^37^, a machine learning model that integrates long-range sequence information to predict gene expression and other histone modifications. Crucially, Enformer only uses sequence information to make predictions, allowing us to compare predictions at genomic locations with and without integrated insulator constructs. Since Enformer does not directly predict gene expression, we focused on predicting Cap Analysis of Gene Expression (CAGE) and H3K27me3 levels. Enformer predicted new CAGE peaks at locations with integrated reporter genes (Supplementary Fig 7A) and higher CAGE fold-changes (insulator/uninsulated) in ALOXE3-up and A2-up compared to unchanged locations (Supplementary Fig 7B), so we focused on these two insulators. In the regions flanking ALOXE3-up locations, Enformer predicted higher levels of H3K27me3 than ALOXE3-unchanged locations (Supplementary Fig 7C), which agrees with our observation that ALOXE3-up are enriched in H3K27me3 chromatin.

At the uninsulated locations, we expect that H3K27me3 from flanking chromatin spreads into the reporter gene to silence expression, resulting in a correlation between H3K27me3 levels in flanking and reporter chromatin. If the insulators block H3K27me3 spreading, then we would expect a decrease in correlation between H3K27me3 levels in flanking and reporter chromatin predicted by Enformer. We find that the correlation between flanking and reporter chromatin was high in the uninsulated and A2 locations, but decreased in ALOXE3 locations, suggesting that only ALOXE3 can block the spread of H3K27me3 (Supplementary Fig 7D). To test whether ALOXE3-up locations are specifically contributing to the decrease in correlation, we compared the correlations between flanking and reporter gene chromatin for ALOXE3-unchanged vs ALOXE3-up locations. ALOXE3-up locations tend to have lower H3K27me3 than expected compared to ALOXE3-unchanged locations (Supplementary Fig 7E). In contrast, A2-up locations and A2-unchanged locations have similar correlations (Supplementary Fig 7F). These results support the idea that ALOXE3 blocks H3K27me3 spreading from flanking chromatin into the reporter genes, leading to upregulation in ALOXE3-up locations.

### cHS4 and A2 enhance expression in ‘primed’ genomic environments

We observed that unlike ALOXE3-up locations, both A2-up and cHS4-up locations seem to be interacting with more enhancers and transcribed regions (Figs 5B & C). This suggested that the non-insulated reporters may be more highly expressed in A2-up and cHS4-up locations. Instead, we found that the expression levels of the non-insulated reporters are generally lower in A2-up and cHS4-up locations compared to the unchanged locations, albeit marginally higher than ALOXE3-up locations (Fig 5D). Thus, we speculate that A2-up and cHS4-up locations are primed for high expression and that A2 and cHS4 act as ‘enhancers’ at these locations by recruiting additional TFs that upregulate expression. This model can also explain the discrepancies between the chromatin environments of A2-up and cHS4-up locations (Fig 5A). While A2 requires a more highly active chromatin environment to act as an enhancer, cHS4 might upregulate expression at more neutral environments because it contains many more motifs and may recruit many more TFs (Supplementary Fig 5G). Consistently, mutant-up locations are all enriched for active histone modifications and looped to more enhancers/transcribed regions (Supplementary Fig 8A-9C). Despite that, the uninsulated reporters are also not highly expressed at mutant-up locations (Supplementary Fig 8D). This suggests that deleting the B-box motifs in ALOXE3 abolishes its ability to act as a heterochromatin barrier.

Finally, there are also a small number of locations (86) that are upregulated by all three insulators. These locations are enriched for both active (particularly H3K4me2 and me3) and inactive histone modifications (Fig 5E), which could represent a poised chromatin environment. In these environments, ALOXE3 can block heterochromatin-mediated silencing, while cHS4 and A2 can also gain enhancer activity to upregulate gene expression. Taken together, our results suggest that only ALOXE3 acts as a classical heterochromatin blocker, while the other constructs appear to act as enhancers at primed locations across the genome (Fig 5F).

## Discussion

Insulators are a class of *cis*-regulatory sequences that have been associated with several functions. Due to the lack of functional assays to measure insulator activity in the genome, it has been difficult to understand or predict insulator function. Here we developed a high-throughput genome-wide assay, MPIRE, to systematically measure the impact of insulators on gene expression at thousands of locations across the genome and show that insulators have distinct, pleiotropic activities depending on their location in the genome. We propose that our results reconcile the differences between genome-wide analyses and analyses of individual insulators. Depletion of insulator-binding proteins such as CTCF or Cohesin have minimal changes on gene expression, consistent with our results that insulators only function in small numbers of genomic locations. Where the insulators do function, different insulators preferentially block specific enhancers that recruit different complements of TFs. Among the three tested insulators, only ALOXE3 protects against heterochromatin silencing. These results explain why one-at-a-time experiments can sometimes show that insulators have large and specific effects on gene expression.

A long-standing question in gene regulation is how enhancers contact and activate target genes in the genome. Recent evidence has shown that enhancers and promoters are broadly compatible with each other, raising the question of how specific interactions are achieved^27,38,39^. Insulators are thought to facilitate this process by blocking enhancers from interacting with their target promoters, but it remains unclear whether insulators are modular elements that can block all enhancers in the genome. Our results show that insulators block enhancers at specific locations in the genome, thereby leading to a reduction in gene expression at those locations (Fig 4G). For example, ADNP binding is enriched in enhancers blocked by ALOXE3 (Fig 4F), and has previously been implicated in modulating 3D genome interactions by competing with CTCF for binding at SINE B2 transposable elements^40^. Both SINE B2 and ALOXE3 elements are transcribed by RNA Pol III and have been shown to act as insulators^20,21,41^, which may explain ALOXE3’s ability to block ADNP-bound enhancers. RNA Pol III has also been suggested to aid ADNP in recruiting distal CTCF sites, which may help ALOXE3 act as an insulator in ALOXE3-down locations^42^. The two insulators that bind CTCF (A2, cHS4) are not any more similar compared to ALOXE3, implying that CTCF itself is not sufficient to explain insulator activity. The specificity of different insulators for different enhancers suggests a model where insulators interact with select enhancers to modulate specific enhancer-promoter interactions.

While we only tested three insulators in this study, our results demonstrate that not all insulators possess barrier activity against heterochromatin. Only ALOXE3 appears to be able to block reporter gene silencing. A parsimonious explanation for this difference might be that ALOXE3 binds to RNA PolIII and gets transcribed^21^, allowing it to generate an open chromatin environment that can overcome heterochromatin spreading. Indeed, the recruitment of transcriptional activators can insulate against repression^43,44^, and transcription of various TEs have been shown to be sufficient to generate a boundary in mammalian cells^45,46^. Other groups have also observed that cHS4 is not sufficient to protect against silencing at all genomic locations^12,18^, and can block HDAC-mediated but not KRAB-mediated silencing^47^. We find instead that cHS4 and A2 can act as enhancers in primed environments to upregulate gene expression (Fig 5F). Neither cHS4 nor A2 show enhancer activity in plasmid reporter assays^9,31^ (Fig 2B), likely because plasmids do not provide the necessary primed environment. Our results are consistent with the previously proposed model that insulation is a dynamic contest between activation and repression^48^. ALOXE3 is transcribed and has the highest activation rate, allowing it to overcome repressed heterochromatin, while A2 has the lowest activation rate and can only overcome very weakly repressive environments. Moving forward, screening and testing more insulators across more locations in the genome will allow us to build a comprehensive and predictive model of insulator activity. These results will also be useful to improve synthetic biology and gene therapy applications where high levels of sustained gene expression are required.

We developed MPIRE to study insulator activity, but we envision that MPIRE can be used for many other questions involving genomic or chromatin context, for example, to study the interactions between TF binding and chromatin or splicing and chromatin or to assay allelic variants in the same genomic contexts. MPIRE can also be coupled with other DNA-based readouts such as Cut&Tag or 4C to measure other aspects of genome regulation in addition to expression. Understanding how the different processes in the genome interact with each other will be important for determining and quantifying their contributions to gene expression regulation.

## Methods

### Landing pad library cloning

All primers and oligos used in this study can be found in Supplementary Table 5. We first cloned each landing pad construct into a Sleeping Beauty (SB) transposon plasmid containing SB ITRs (gift from Robi Mitra lab). Each landing pad consists of a hsp68 promoter driving the expression the mEmerald reporter gene and is flanked with φC31 and BxbI attP sites for recombination. The φC31 attP site, hsp68 promoter, mEmerald fluorophore, poly(A) signal constructs were amplified by PCR from different plasmids and assembled using the NEBuilder HiFi DNA Assembly Master Mix (HiFi Assembly, NEB #E2621). The BxbI attP site was then added using the Q5 Site-Directed Mutagenesis Kit (Q5 SDM, NEB #E0554). To add a library of diverse random barcodes, we first digested the landing pad plasmid with XbaI at 37℃ for 16 hours. We then ordered an oligo containing 16 Ns with flanking homology arms to the landing pad plasmid (GWLP P1) and used HiFi Assembly (NEB #E2621) to assemble the oligo to the plasmid (50℃, 15 min).

### Generation of cell lines

K562 cells were maintained in Iscove’s Modified Dulbecco′s Medium (IMDM) + 10% FBS + 1% non-essential amino acids + 1% penicillin/streptomycin. To generate pools of cells containing landing pads, K562 cells were electroporated with the SB100x transposase (gift from Robi Mitra lab) and landing pad library at a 1:1 ratio. Specifically, we electroporated 4.8 million cells with 20μg SB100x and 20μg landing pad library with the Neon Transfection System 100μL Kit (Life Technologies, #MPK10025). The cells were cultured for a week then sorted into pools of 2000 cells to bottleneck the numbers of gBCs per pool, ensuring that each gBC is associated with a unique location in the cell^23^. In total we generated 6 pools of cells, with pools 1-2 from the same initial transfection pool (experiment v1) and pools 3-6 from a second initial transfected pool of cells (experiment v2).

To measure expression of each landing pad we harvested RNA and genomic DNA for each pool of cells using TRIzol reagent (Invitrogen #15596026). 4μg RNA was treated with DNAse using the Rigorous DNase treatment procedure from the Turbo DNase protocol (Invitrogen #AM2238) and reverse transcribed with the SuperScript IV First Strand Synthesis System (Invitrogen #18091050). Barcodes were amplified from the library using Q5 High-Fidelity 2X Master Mix (Q5, NEB #M0492) using primers GWLP P2-3 with 20 cycles. We performed 32 PCR reactions per cDNA sample. For genomic DNA, we performed 4 PCR reactions with 1μg DNA per PCR. PCRs from the same samples were then pooled and purified. Sequencing adapters were then added with two rounds of PCR with Q5 (NEB #M0492) (GWLP P4-7). The resulting libraries were sequenced on the Illumina NextSeq platform.

To map the locations of each landing pad we followed the protocol for mapping TRIP integrations as described in our previous study^27^. The resulting libraries were sequenced on either the Illumina NextSeq or NovaSeq platform.

### MPIRE insulator library construction

We first generated a transfer vector containing a reporter gene (hsp68 promoter + mScarlet) flanked by φC31 and BxbI attB sites using HiFi Assembly (NEB #E2621). The insulator sequences were obtained from their respective papers (cHS4 from Chung *et al*.^9^, A2 from Liu *et al*.^31^ and ALOXE from Raab *et al*.^21^). For ALOXE3, we selected to use the sequence with 2 tDNAs rather than 4 tDNAs because it is a tractable length for cloning (∼1kb), does not contain a CTCF binding site and is sufficient for enhancer-blocking activity^21^. cHS4 and A2 were synthesized as gBlocks (Integrated DNA Technologies) and homology arms for the backbone were added by PCR with GWLP P8-9 (cHS4) or GWLP P10-11 (A2). ALOXE3 was amplified from K562 genomic DNA using primers GWLP P12-13 and homology arms were added by PCR with GWLP P14-15. The constructs were then inserted between the φC31 attB site and hsp68 promoter in the transfer vector by HiFi Assembly (NEB #E2621). The cHS4 mutant design is from Farrell *et al*.^32^ (cHS4 x3) and was introduced to the cHS4 transfer vector with Q5 SDM (NEB #E0554) using primers GWLP P16-17. While the CTCF binding site in A2 has been identified, it has not been tested for its dependence on CTCF. Thus, we used the CTCF PWM (JASPAR^49^ MA0139.1) to identify and mutate the three most informative sites in the motif (<u>C</u>ACCA<u>G</u>GT<u>G</u>GCGCT → <u>T</u>ACCA<u>C</u>GT<u>T</u>GCGCT). The mutation was introduced to the A2 transfer vector with Q5 SDM ((NEB #E0554) using primers GWLP P18-19. For ALOXE3, the B-box motifs were identified using the B-box frequency matrix from Pavesi *et al*.^50^ with FIMO^51^, and both motifs were deleted with Q5 SDM (NEB #E0554) using primers GWLP P20-23. Finally, the A2 sequence was randomly scrambled to generate A2 scrambled and synthesized with homology arms as a gBlock (Integrated DNA Technologies), then inserted between the φC31 attB site and hsp68 promoter in the transfer vector by HiFi Assembly (NEB #E2621). Insulator barcodes were then added to each construct by site-directed mutagenesis. All insulator sequences and mutants can be found in Supplementary Table 6.

To test the efficiency of recombination, we generated constructs barcoded with a diverse library of random barcodes. We digested the no insulator and A2 constructs with NheI at 37℃ for 16 hours. We then ordered an oligo containing 16 Ns with flanking homology arms to the plasmid (GWLP P24) and used NEBuilder HiFi DNA Assembly to assemble the oligo to the plasmid (50℃, 15 min).

### MPIRE cell pool validation

For the pilot experiments we focused on cell pool 1. To determine whether the landing pads would recombine with each other, we cotransfected 2.4 million pool 1 cells with 10μg φC31 integrase and 10μg BxbI integrase as described above. The cells were then allowed to grow for a week, and RNA and DNA was harvested using TRIzol reagent (Invitrogen #15596026). We then amplified barcodes from both RNA and DNA to measure expression and mapped the barcodes as described above in the Generation of cell lines section.

To determine the efficiency of recombination we focused on pools 1 and 2. For each pool, we cotransfected 2.4 million cells with a mix of 2.5μg φC31 integrase, 2.5μg BxbI integrase and 5μg of either the no insulator or A2 construct as described above. The cells were then allowed to grow for a week, and RNA and DNA was harvested using TRIzol reagent (Invitrogen #15596026). The RNA was treated with DNase and reverse transcribed to generate cDNA as described above. We then amplified barcodes from both cDNA and DNA in 4 PCR reactions per sample using Q5 (NEB #M0492) and primers specific to the reporter genes (GWLP P3, P25). We pooled PCRs from the same samples for PCR purification, then used 4ng of the product for further amplification with two rounds of PCR to add Illumina sequencing adapters (GWLP P3, P33, P6-7). The resulting libraries were sequenced on the Illumina NextSeq platform.

### Insulator plasmid activity

To measure expression from plasmids alone, we transfected K562 wild-type cells with each construct separately. For each construct, we performed three biological replicates. We transfected 1.2 million cells with 5μg plasmid DNA per replicate using the Neon Transfection System 100 μL Kit (Life Technologies #MPK10025). RNA was harvested 72 hours after transfection using the Monarch Total RNA Miniprep Kit (NEB #T2010). mScarlet and HPRT (housekeeping gene for normalization) levels were then measured by qPCR using the Luna Universal One-Step RT-qPCR Kit (NEB #E3005) with 100ng total RNA per reaction, three reactions per replicate (mScarlet primers: GWLP P25-26, HPRT primers: GWLP P27-28). Expression was calculated using the ΔΔCt method, normalized to the no insulator construct.

### Enhancer-blocking assay

For the enhancer-blocking assay, we designed plasmids that contain either enhancer-promoter or enhancer-insulator-promoter sequences to test whether the insulator can block enhancer-mediated activation of promoter activity. We first selected an enhancer that was previously shown to be active with the hsp68 promoter in K562 cells^52^. The full sequence of the enhancer is: GCCCCCCTTCTTCCTATGTCTGATGGAGTTTCCTCTCTAAGTAGCCATTTTATTCTGCTGACTCACCCTCTAACTCCCGGTCTTATTCCATCCTGCCTCAGGGTCTGTGGTGTAGTCATAGCACATGCATCTCCTCCGGCTCGCTGATT

The no insulator construct was digested with EcoRI to insert the enhancer upstream of the hsp68 promoter. The enhancer was amplified by PCR to add overhangs to the backbone vector (GWLP P29-30) and assembled into the backbone by HiFi Assembly (NEB #E2621).

To add the different insulator constructs, the +enhancer plasmid was digested with AgeI and EcoRI. For A2, A2-mut, A2-scrambled, cHS4 and cHS4-mut, we digested the respective plasmids with AgeI and EcoRI and ligated them to the backbone with T4 DNA Ligase (NEB #M0202). For ALOXE3 and ALOXE3-mut, we amplified the insulators by PCR to add overhangs to the backbone vector and assembled the insulators into the backbone by HiFi Assembly (GWLP P31-32).

We measured expression from these constructs in the same way as described in the Insulator Plasmid Activity section, except with two biological replicates instead of three per construct.

### MPIRE

We pooled insulator constructs for transfection into landing pad cell pools. In the first experiment, we combined the no insulator, A2, A2 mut, A2 scrambled, cHS4, and ALOXE3 constructs and tested them in pools 1 and 2. In the second experiment, all the constructs were combined in equal amounts and tested in pools 3-6. We used the Neon Transfection System 100μL Kit (Invitrogen #MPK10025) for transfection. Each pool was transfected separately and we performed two replicates per pool. For each replicate, we performed four transfections, each with 1.2 million cells and a mix of 2μg φC31 integrase, 2μg BxbI integrase and 4μg of the insulator constructs pooled in equal ratios.

We harvested DNA and RNA from cells one week after transfection using the TRIzol Reagent (Invitrogen #15596018). The RNA was treated with DNase and reverse transcribed to generate cDNA as described above. Barcodes were amplified from cDNA using Q5 (NEB #M0492) with primers specific to the reporter gene (GWLP P3, P25), with 32 PCR reactions per biological replicate. Similarly, barcodes were amplified from genomic DNA with 8 PCR reactions per biological replicate. We pooled PCRs from the same biological replicates for PCR purification, then used 4ng of the product for further amplification with two rounds of PCR to add Illumina sequencing adapters (GWLP P4, P33, P6-7). The resulting libraries were sequenced on the Illumina NextSeq platform.

### Data analysis

The preprocessing of sequencing reads was performed with Python 3.9. All statistical analyses and figures were done in R 4.2.0. We used bedtools v2.30.0 for genomics analyses.

### MPIRE cell pools data processing

For expression from the initial cell pools and cells transfected with integrase only, we first filtered for reads that contained the gBC in the correct sequence context. We then filtered for gBCs with >20 reads and normalized the reads by sequencing depth. We then calculated expression as log_2_(RNA counts/DNA counts) for all barcodes. The expression of the landing pads in the original cell pool and in the integrase-transfected cells can be found in Supplementary Tables 2 and 7, respectively.

To identify the locations of the landing pads, we obtained paired-end reads containing the barcode on one read and the sequence of the integration site on the other. We matched the barcodes with the integration site sequence, then aligned them to hg38 with BWA using default parameters. We only kept barcodes where the reads for one location represented at least 80% of all the locations for that barcode. The mapped integration locations can be found in Supplementary Table 1. For pools from the same initial transfection (Pools 1&2, Pools 3-6), a small number of barcodes are shared between them because the same cells were sorted into multiple pools. These barcodes are indicated with Px instead of a pool number. To compare locations before and after integrase-only transfection, we compared the mapped locations in pool 1, only considering gBCs with exact matches. These locations can be found in Supplementary Table 8.

### Recombinase efficiency data processing

To evaluate the efficiency of recombination, we first filtered for reads that contained both barcodes (rBC and gBC) in the correct sequence context. We then filtered for barcode pairs that had >20 reads associated with them and tabulated the number of rBCs/gBC for either the no insulator or A2 construct (Supplementary Table 9). For the proportion of recovered LPs, we compared the DNA gBCs recovered from cells transfected with either the no insulator or A2 construct to the DNA gBCs recovered from the original cell pools.

### MPIRE data processing

For the recombined MPIRE libraries, we first filtered for reads that contained both barcodes (iBC and gBC) in the correct sequence context. The gBCs were then compared to the list of mapped gBCs (gBCs that have a unique location associated with them, Supplementary Table 1). A gBC was assigned to a mapped gBC if both barcodes have a hamming distance <5 and the gBC is ≥5 hamming distance from all other gBCs (the average hamming distance between mapped gBCs is 9). We then tabulated the number of barcodes or barcode pairs after assigning gBCs to a mapped gBC for both RNA and DNA.

To calculate reproducibility and to compare expression before and after recombination, we first filtered for gBC-iBC pairs that had >20 reads and normalized the reads by sequencing depth. We then calculated expression as log_2_(RNA counts/DNA counts) for all barcodes that are present in both the RNA and DNA pools. The calculated expression values can be found in Supplementary Table 10.

To calculate expression after recombination of the insulator library, including those that might not have an RNA count because they are lowly expressed, we turned to the MPRAnalyze tool^53^. We added a pseudocount of 1 to all counts and used the analyzeQuantification function to calculate the transcription rate (α) for each construct at each location using counts from both biological replicates. If the same barcode is present in multiple pools, then the counts from all replicates across the pools were used as input, so that each location only gets one α value. Expression values can be found in Supplementary Table 3. To quantify the effects of the insulators, we used the analyzeComparative function in MPRAnalyze to calculate the fold-change of the insulated vs uninsulated locations. The output of this analysis can be found in Supplementary Table 4.

### Histone modifications

We downloaded ChIP data for various histone modifications in K562 from the ENCODE database^54^ (Supplementary Table 11). We used bedtools^55^ to map the mean histone signals in the 20kb surrounding each landing pad (10kb upstream and downstream). For each histone modification, we standardized the signal values, then computed the average signal for ins-unchanged, ins-up and ins-down locations respectively. All heatmaps were generated using the ComplexHeatmap package in R^56^.

### Other genomics analysis

The core 15-state chromHMM annotation for K562 cells was downloaded from the Roadmap Epigenomics Project^57^ and similar annotations were grouped. We overlapped the landing pads with the corresponding annotation using the GenomicRanges R package^58^.

ATAC-seq data for K562 cells was downloaded from the ENCODE database^54^ (Supplementary Table 11). We used bedtools^55^ to count the number of ATAC-seq peaks around each landing pad.

To identify locations that are looped to each location, we used K562 Hi-C data that was previously generated and processed^59^. We then used the peakHiC tool^60^ to call loops for each landing pad with the following parameters: window size = 80, alphaFDR = 0.5, minimum distance = 10kb, qWr = 1. The regions identified by peakHiC were overlapped with chromHMM annotations using the GenomicRanges R package^58^ to identify putatively interacting enhancers with each landing pad. All locations looped to each landing pad can be found in Supplementary Table 12.

### Transcription factors enrichment analysis

We downloaded all available TF ChIP data in K562 from the ENCODE database^54^. The full list of downloaded files can be found in Supplementary Table 13. For this analysis we focused on the locations that are downregulated by only one or two insulators, locations that were downregulated by all three were excluded. We used bedtools^55^ to count the number of TF ChIP peaks in the surrounding 20kb of each landing pad (10kb upstream and downstream) or to count the number of peaks in each putative enhancer identified above. We compared the number of peaks between different groups of landing pads or enhancers using chi-squared tests followed by *p*-value correction with the Benjamini & Hochberg (BH) method.

### Enformer predictions

We decided to use Enformer because it can predict histone modifications and expression directly from sequence^37^. We first downloaded the published, trained model from TensorFlow Hub (https://tfhub.dev/deepmind/enformer/1). For each landing pad, we considered the landing pad to be the center of the sequence to use as input and extended it out on both flanks using the hg38 genome for a total length of 393,216bp, which was the length used in the training model. The model predicts signals for the central 114,688bp, which includes the landing pad. For each location, we used the model to predict CAGE and H3K27me3 in K562 cells. To calculate CAGE signals, we summed CAGE scores across all windows that contain the landing pad. We then calculated fold-changes for each insulator by taking log_2_(predicted_insulator_CAGE/predicted_uninsulated_ CAGE) (Supplementary Table 14). To calculate H3K27me3 signals, we either summed across all windows that contain the landing to get the landing pad H3K27me3 signals, or we summed across all windows outside the landing pad (that were used in the input data) to get flanking H3K27me3 signals (Supplementary Table 15). The values were normalized by z-scoring the predicted values for each insulator.

## Acknowledgements

We are grateful to Robi Mitra for providing us with plasmids for our experiments. We thank members of the Cohen Lab for helpful discussion and critical feedback on the manuscript. We also thank Jessica Hoisington-Lopez and MariaLynn Crosby in the DNA Sequencing Innovation Lab for assistance with high-throughput sequencing and the Genome Engineering and iPSC Center for kindly allowing us to use their flow cytometer for cell sorting. This work was supported by grants to BAC from the National Institutes of Health (R01GM092910).

## Author contributions

C.K.Y.H and B.A.C conceived and designed the project. C.K.Y.H, A.A.E and J.L conducted the experiments, C.K.Y.H, A.A.E and A.J.F performed the analyses. C.K.Y.H and B.A.C wrote the manuscript.

## Supplementary Figures

**Supplementary Fig 1:**
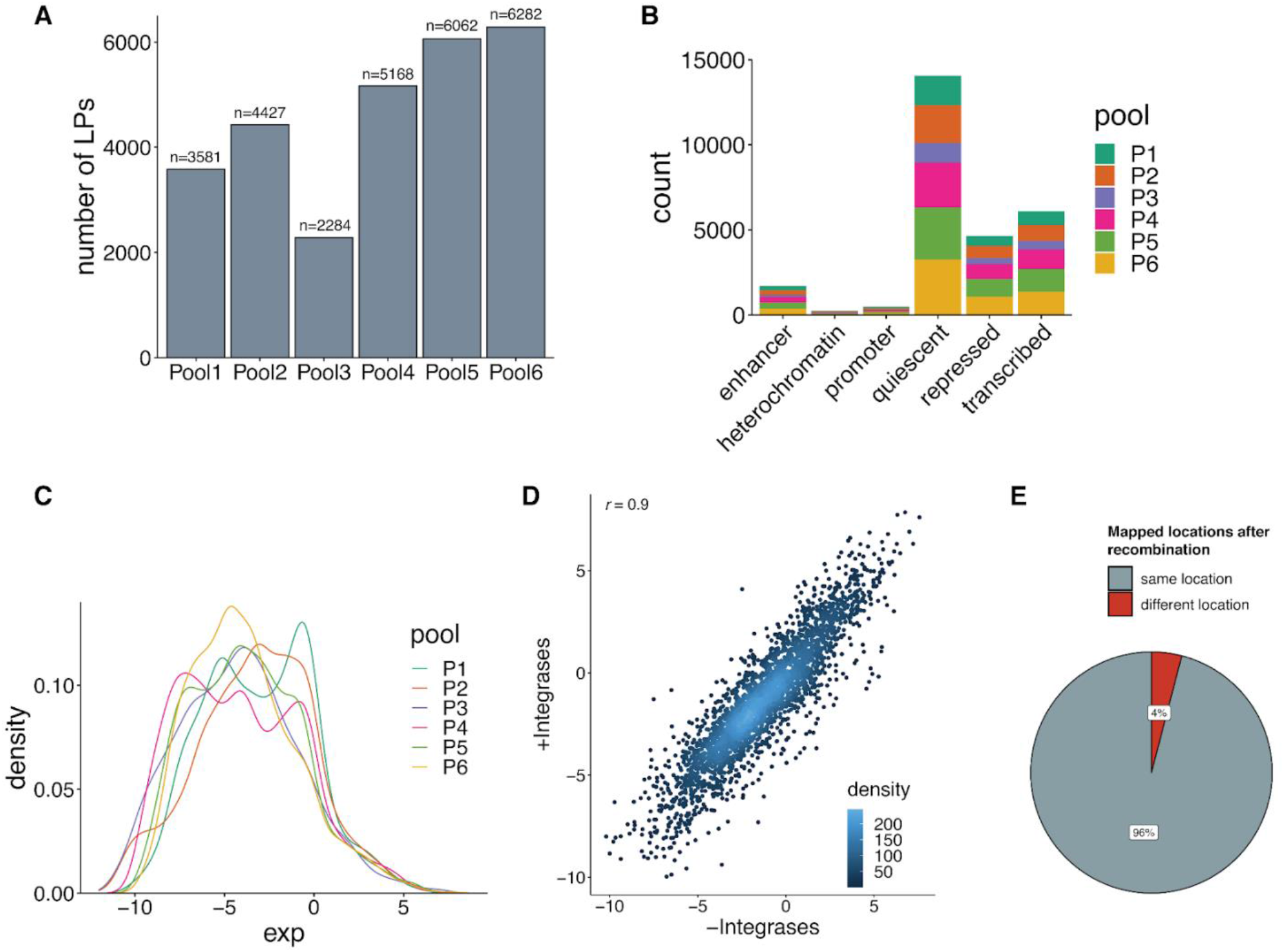
Overview of cell pools for MPIRE. **(A)** Barplot of the number of mapped landing pads per pool. The exact number is indicated above each bar. **(B)** Barplot of the number of landing pads in each chromHMM type per pool. **(C)** Expression across all landing pads in each pool. **(D)** Expression of landing pads before and after transfection with φC31 and BxbI integrases only. Correlation indicated is Pearson’s *r*. **(E)** Percentage of locations that mapped to the same or different locations after transfection with the integrases only.

**Supplementary Fig 2:**
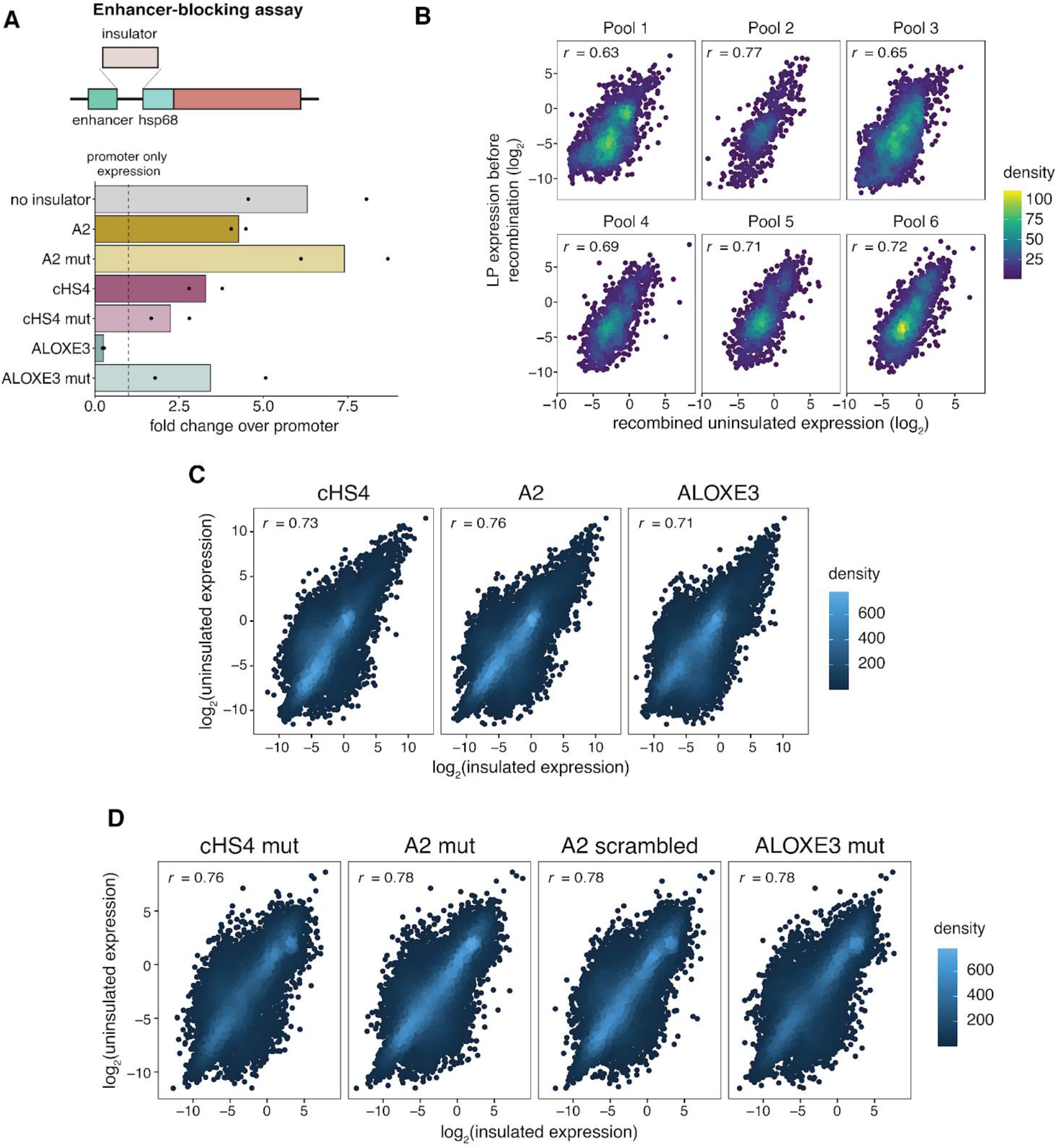
MPIRE reliably measures insulator activity. **(A)** Enhancer-blocking assay for testing insulator activity. Expression was measured by qPCR and normalized to the promoter-only (hsp68) construct. Each dot represents a measurement from one replicate, two transfection replicates were performed for each construct. **(B)** Correlation between reporter gene measurements from the initial landing pad and from the recombined uninsulated construct for each pool. **(C)** Correlation between uninsulated construct and respective insulator at each location. **(D)** Correlation between uninsulated construct and respective mutant construct at each location. Correlations shown are Pearson’s *r*.

**Supplementary Fig 3:**
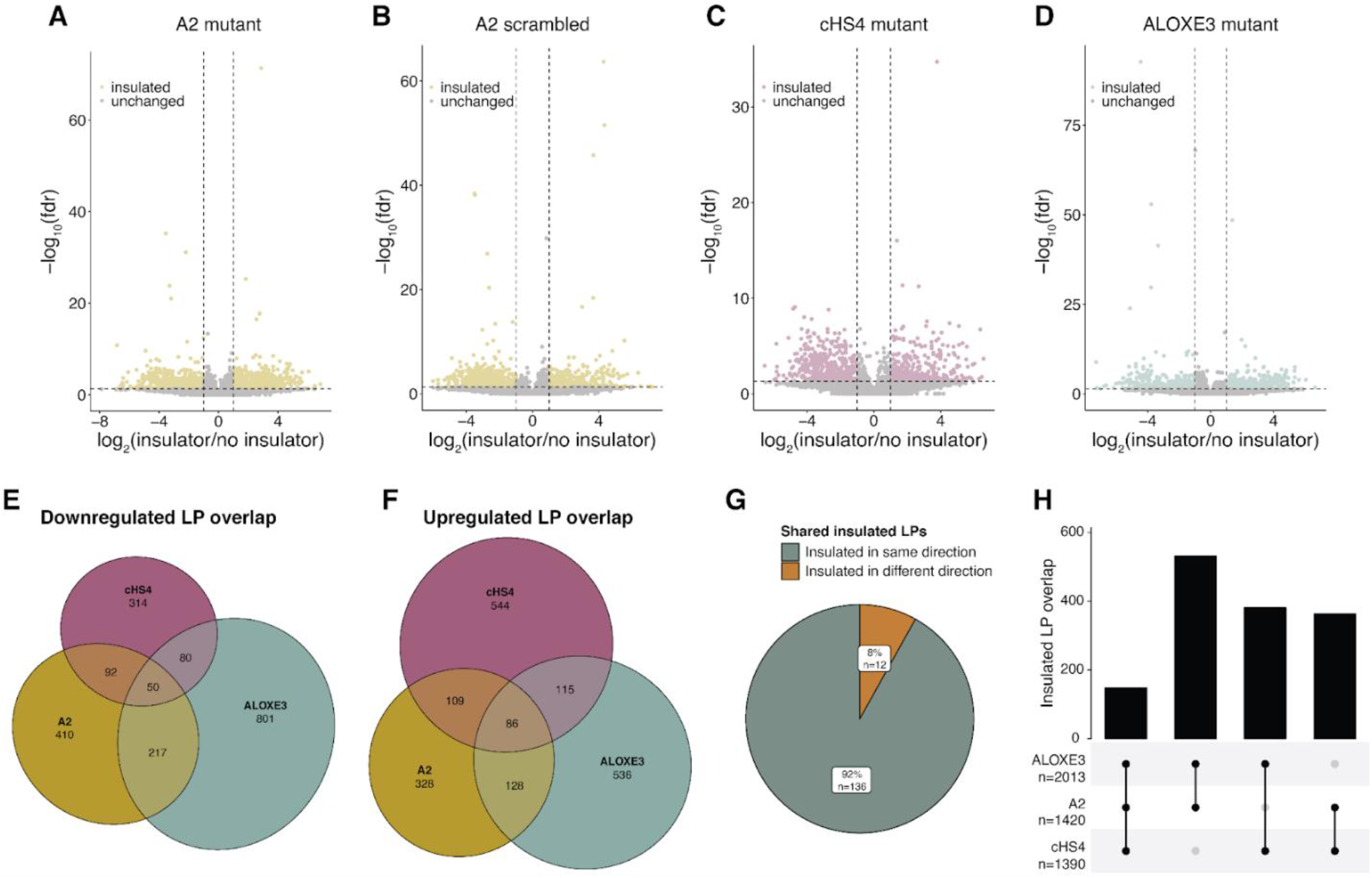
Insulators function at distinct locations in the genome. **(A-D)** Volcano plot of insulator effect across locations for A2 mutant (n=14390) **(A)**, A2 scrambled (n=13690) **(B)**, cHS4 mutant (10944) **(C)** and ALOXE3 mutant (n=11135) **(D)**. Colored points (insulated locations) represent locations where log_2_(insulator/no insulator) > 2 and fdr < 0.05. **(E)** Overlap between locations insulator-down by the respective insulators. **(F)** Overlap between locations upregulated by the respective insulators. **(G)** Percentage of shared insulated locations (insulated by all insulators) that are insulated in the same or different direction. **(H)** Upset plot of the number of shared insulated locations by the indicated pairs.

**Supplementary Fig 4:**
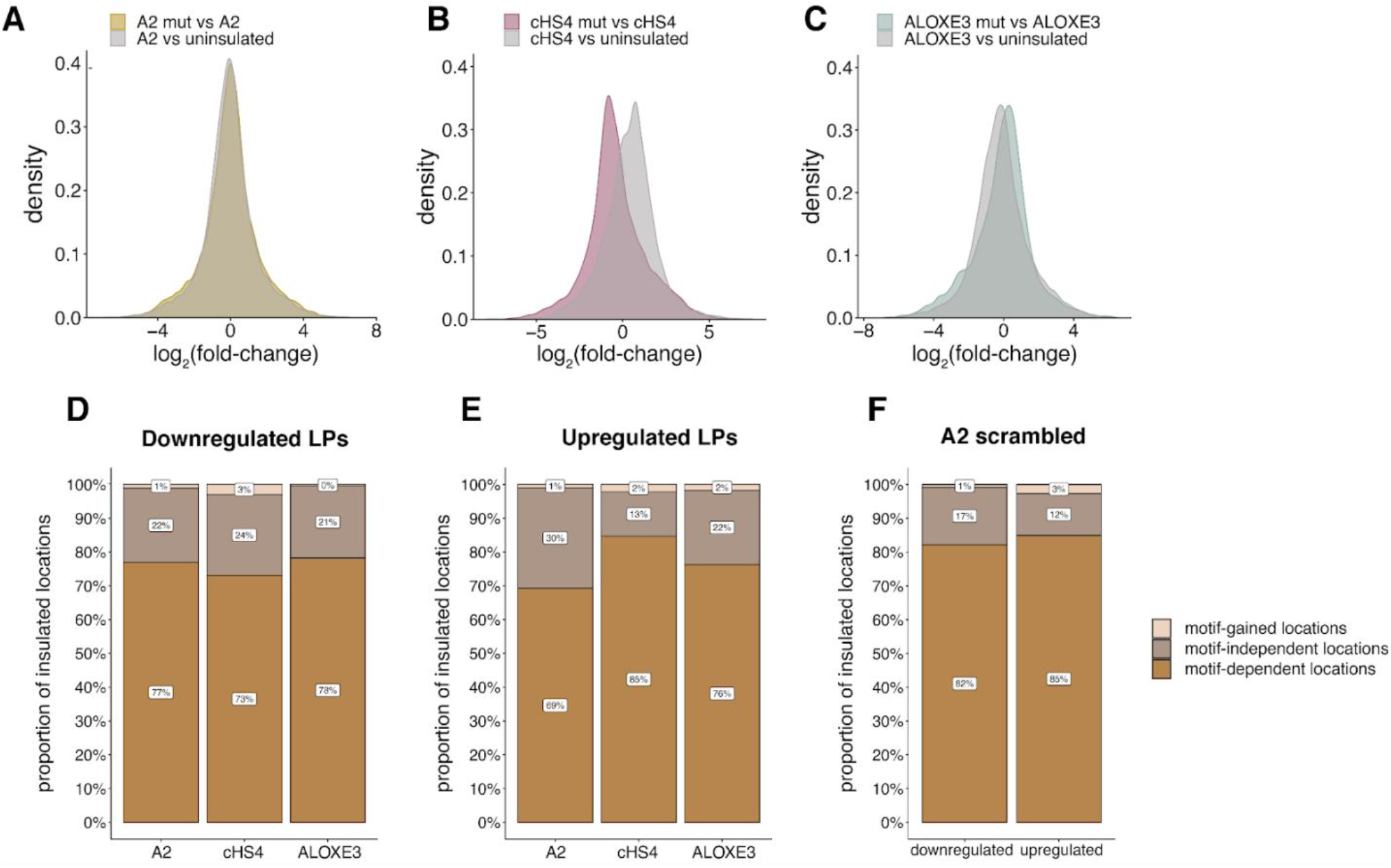
Insulators are mostly dependent on their respective sequence motifs. **(A-C)** Fold-changes between either the insulator and uninsulated constructs or insulator and its mutant construct for A2 **(A)**, cHS4 **(B)** or ALOXE3 **(C). (D, E)** Percentage of insulated locations that are motif-gained, motif-independent or motif-dependent for insulator-down (D) or insulator-up (E) locations. **(F)** Percentage of insulated locations in each category for A2-scrambled-up and A2-scrambled-down locations.

**Supplementary Fig 5:**
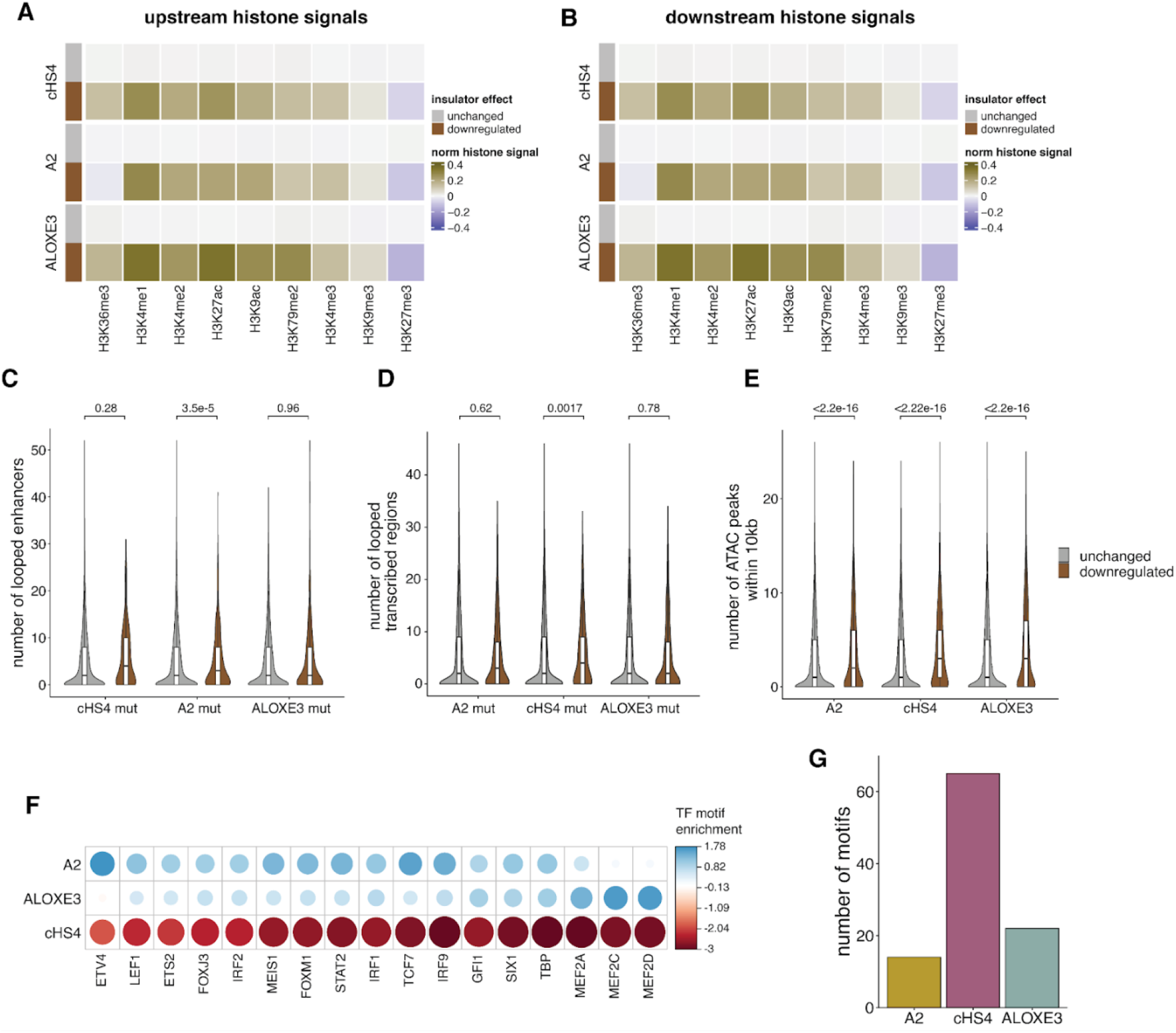
Insulators block specific enhancers in the genome. **(A, B)** Mean normalized histone modifications either 10kb upstream (A) or downstream (B) of insulator-unchanged and insulator-down location. Histone signals are standardized across locations for each modification. **(C)** Violin plot of number of looped enhancers per location for mutant-unchanged vs mutant-down locations. **(D)** Violin plot of number of looped transcribed regions per location for mutant-unchanged vs mutant-down locations. **(E)** Violin plot of the number of ATAC peaks for insulator-unchanged vs insulator-down locations. *p*-values (two-sided Wilcoxon test) are shown above each plot. **(F)** For all insulators, enhancers that are looped that the insulator-down locations were compared for differences in TF motif content (Chi-squared test). The residuals of motifs that are significantly different (*p* < 0.05) after Benjamini & Hochberg (BH) correction was plotted. **(G)** Number of TF motifs in each insulator.

**Supplementary Fig 6:**
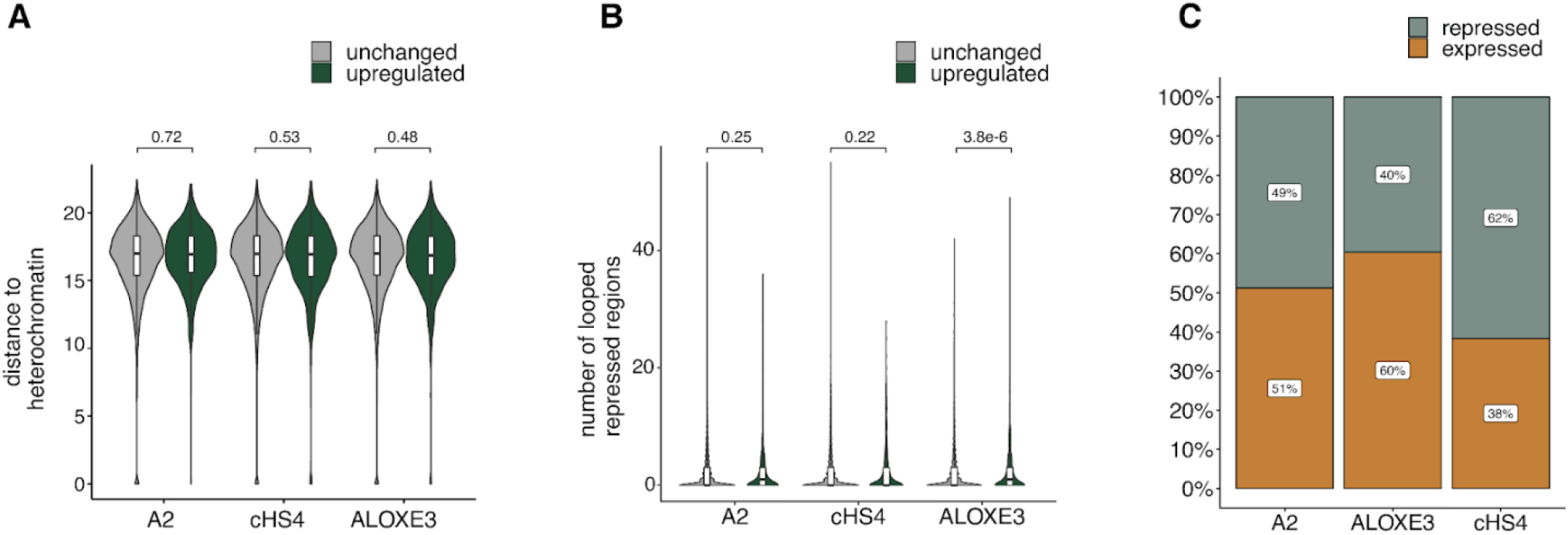
ALOXE3 acts as a barrier against heterochromatin. **(A)** Distance to closest heterochromatic region for each insulator-unchanged and insulator-up location. **(B)** Number of regions looped to each location designated as repressed by chromHMM. *p*-values (two-sided Wilcoxon test) are shown above each plot. **(C)** Proportion of locations that are expressed (contains detectable RNA counts) vs repressed (no detectable RNA counts) for each insulator.

**Supplementary Fig 7:**
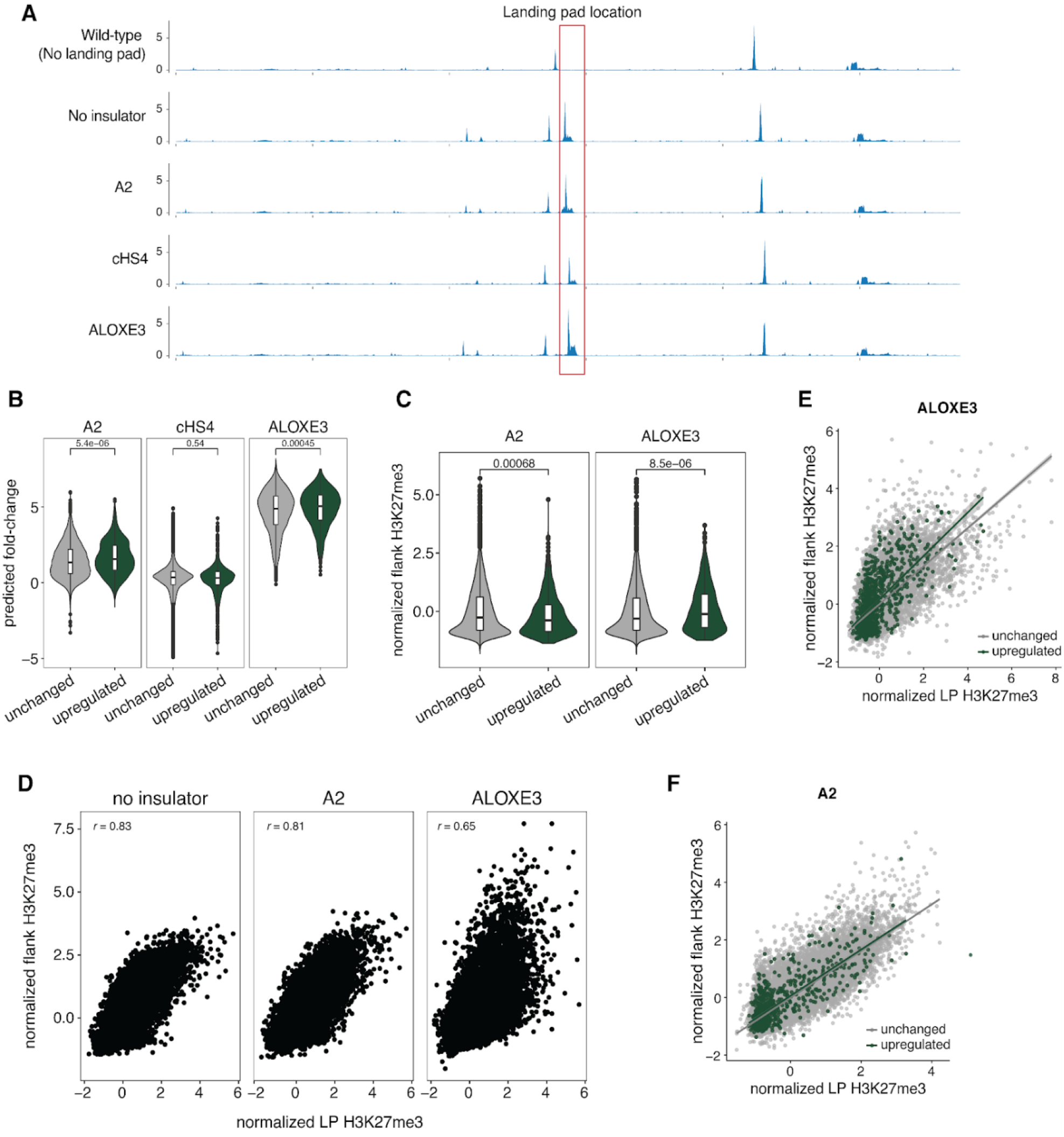
Enformer predicts ALOXE3 barrier activity and loss of H3K27me3. **(A)** Representative example of predicted CAGE signals by Enformer for all insulator constructs. Each track is 114,688bp centered on the site of the landing pad insertion. **(B)** Violin plot of fold-changes in CAGE signal (calculated as log_2_(insulator/uninsulated)) in insulator-unchanged vs insulator-up locations. **(C)** Violin plot of summed H3K27me3 levels in regions flanking the landing pad for insulator-unchanged vs insulator-up locations. *p*-values (two-sided Wilcoxon test) are shown above each plot. **(D)** Correlation between summed H3K27me3 levels in the LP and flanking regions. Correlations shown are Pearson’s *r*. **(E, F)** Same correlation plot in **D** for the ALOXE3 **(E)** and A2 **(F)** constructs but colored by whether the points are insulator-unchanged or insulator-up locations. The line presents the best fit linear model with 95% confidence interval surrounding the line in light gray. All H3K27me3 values are z-scored for normalization.

**Supplementary Fig 8:**
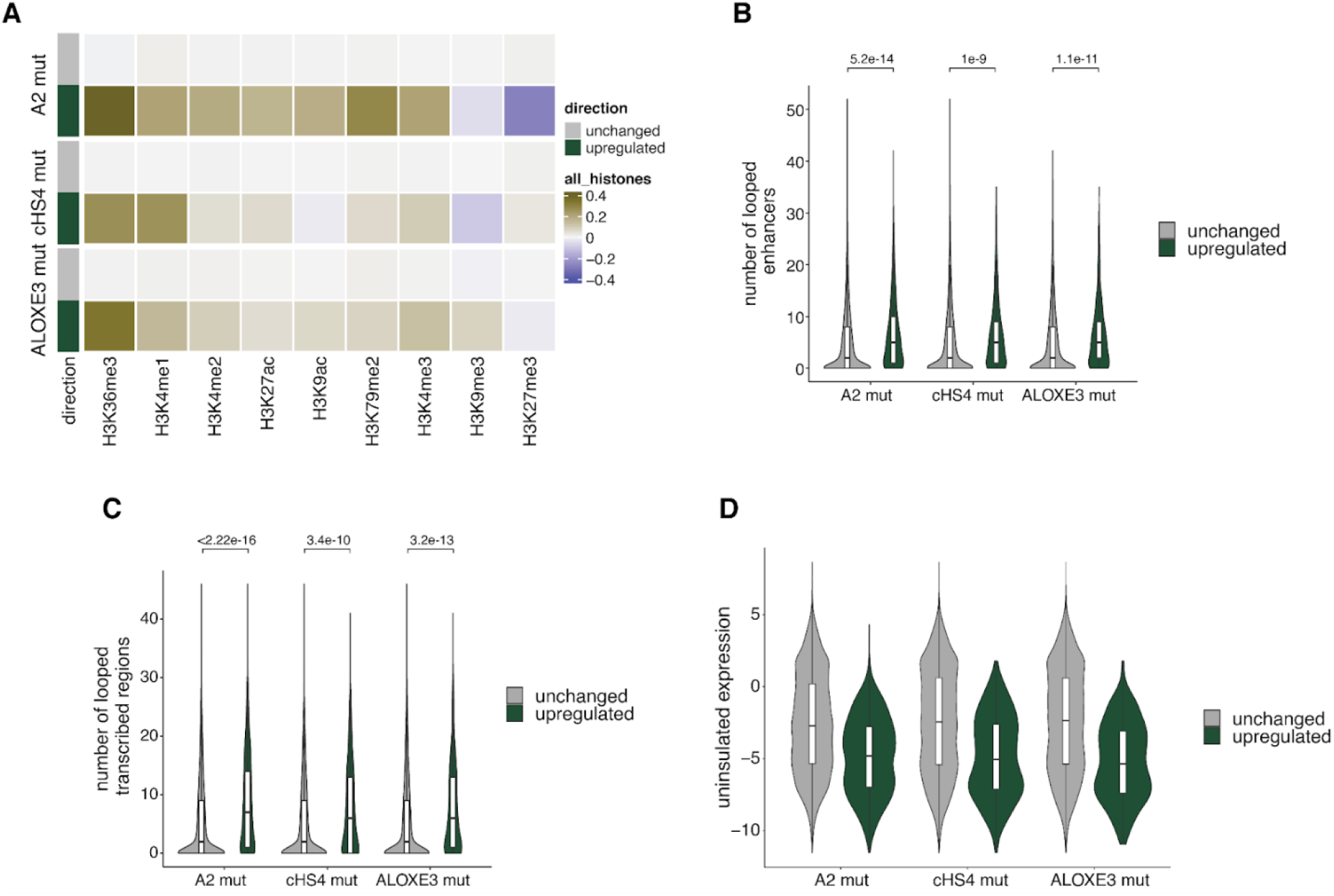
cHS4 and A2 act as enhancers at ‘primed’ genomic locations. **(A)** Mean normalized histone modifications for mutant-unchanged and mutant-down locations. Histone signals are calculated as the mean of 10kb surrounding each location and standardized across locations for each modification. All mutants are enriched for ‘active’ histone modifications. **(B)** Violin plot of number of looped enhancers per location for mutant-unchanged vs mutant-up locations. **(C)** Violin plot of number of looped transcribed regions per location for mutant-unchanged vs mutant-up locations. *p*-values (two-sided Wilcoxon test) are shown above each plot. **(D)** Expression distribution of the uninsulated reporter at mutant-unchanged vs mutant-up locations.

